# A Course-Undergraduate Research Experience (CURE) to explore the effect of structural variants on gene expression in *C. elegans* balancers

**DOI:** 10.64898/2026.01.21.700799

**Authors:** Tatiana Maroilley, Victoria Rodrigues Alves Barbosa, Rumika Mascarenhas, Suzanne Ferris, Catherine Diao, Fatemah AlAwadhi, Saood Aldakheel, Ahmad Ali, Denah Alkanderi, Meshary Alshatti, Samyah Alsuwaileh, Khadeeja Asghar, Ryan Bui, Bentley Chai, Lara Dsouza, Parisa Etemadi Nezhad, Emily Garcia-Volk, Zainub Haq, Sadeeb Hossain, Grace Johnson, Nidhi Kotikalapudi, Inara Lalani, Courtney Lenz, Taye Louie, Stephen Moore, Shubh Patel, Shuvam Prasai, Ronaar Qureshi, Farah Rahmani, Bilal Shakir, Sadhiya Siraj Ahamed, Hong Anh Tran, Radia Waziha, Cassandra M. Wood, Sara Zbinden, David Anderson, Maja Tarailo-Graovac

**Author notes:** Corresponding authors: Maja Tarailo-Graovac and Tatiana Maroilley.

## Abstract

Bioinformatics, a discipline at the crossroads of Biology and Computational Sciences, also referred to as Computational Biology, is nowadays widely spread in research programs. However, implementing any Bioinformatics projects requires the ability to comprehend biological concepts and apply computational approaches, and rare are the undergraduate programs offering such multi-disciplinary training. In addition, understanding the dynamic between Biology research projects and Bioinformatics analyses is challenging with no real-life experience. Course-based undergraduate research experience (CURE) courses are innovative programs that allow more students to acquire research experience and provide the perfect setting to introduce students to applied bioinformatics. As a part of the Bachelor of Health Sciences of the Cumming School of Medicine at the University of Calgary (Canada), a CURE applied bioinformatics was implemented in the Winter of 2023 to 2025. Students investigated the effect of structural variants (SVs, genetic variants larger than 50 bp) on gene expression in the model organism *Caenorhabditis elegans* (a hermaphrodite 1-mm long roundworm). The students detected and characterized SVs by analyzing genome and transcriptome sequencing data of *C. elegans* strains called balancers, as they are known to carry large genomic variations balancing regions of the genome by limiting recombination and allowing maintenance of lethal mutations. They used Galaxy, a public web-based supercomputing resource, but also a local High-Performance computing system, and R, to report different effects of SVs on gene expression and splicing. Students’ research explained the molecular mechanism behind the uncoordinated phenotype caused by the reciprocal translocation eT1(III;V) and uncovered unexpected effects on gene expression on an understudied gene. We evaluated the course’s impact on student learning journeys and showed that the CURE favored students’ understanding of the Bioinformatics field and fostered their research interest. We provide here guidelines to facilitate the CURE implementations to improve access for undergraduate students to bioinformatics research experiences.

## Introduction

For over a decade, innovative teaching methods are encouraged to increase interest for research, improve students’ learning experience, and allow the acquisition of trans-disciplinary skillset [1–7]. In higher education, research-based programs such as Course-based Undergraduate Research Experience (CURE) fostering active student-centered teaching methods are flourishing in different fields of research [8–21], including Bioinformatics [22–28]. CUREs were originally inspired by apprenticeship, adapted to a classroom setting with a large audience [29]. In CUREs, students apply scientific methods to authentic and novel research projects of interest for the scientific community [23]. CUREs are then inquiry-based, which has been shown to improve student performance, their problem-solving and analytical skills, confidence, and interest toward research [12, 23, 30–33]. For Instructors, CUREs are an opportunity to unite teaching and research activities. CUREs also provide an opportunity to develop and implement innovative teaching methods, by reducing the amount of formal lectures, and fostering mentorship activities [34, 35].

Bioinformatics emerged originally as an interdisciplinary research field from the increasing need for Computer Sciences approaches in Life Sciences research, concomitant with the advent of high throughput technologies and development of high-performance computing systems. Nowadays, Bioinformatics, also referred to as Computational Biology or Data Sciences, has been exponentially expanding and requires more and more high-qualified personals. As the field is rapidly evolving, it is then challenging to define simply and clearly, limiting its attractiveness for undergraduate students. In Bioinformatics, Computational Biology, or Data Science, one obstacle to the implementation of CUREs is the challenge for students to have prior pertinent knowledge in both Biology and Computational Sciences. This results in an impression of reduced accessibility for students at registration and causes anxiety for students who enrolled in the class.

In addition, it requires multi-disciplinary training with hands-on experience and knowledgeable instructors, opportunities not always available to students. As we entered the era of omics data, computational biologists must adapt to different datasets (genomics, transcriptomics, metabolomics, proteomics…). But each type of data requires an understanding of the technologies used to acquire the data, and of the relevant molecular mechanisms. In addition, depending on the size of the dataset, mathematical modelling and statistical approaches might be required. With the advent of artificial intelligence, even more advanced computational techniques will become routine in Bioinformatics.

In the field of genomics, the advent of sequencing technologies has greatly improved our ability to explore genome variability. Structural variants (SVs) are defined as variations or mutations in the genome larger than 50 bp [36]. They can be balanced (translocations or inversions) or unbalanced (copy number variants such as deletions or duplications) [37]. In addition, complex genomic rearrangements (CGRs) are defined as an overlapping set of structural variants, impacting one or a few genomic loci. The most complex ones have been defined as chromoanagenesis (including chromothripsis, chromoanasynthesis, and chromoplexy), resulting from a catastrophic series of variations and DNA breakages re-shuffling entirely one or a few genomic regions [38]. If SVs and CGRs were known to be involved in phenotypes, diseases, adaptation and evolution [39], they remained until recently understudied due to the challenges inherent to their detection. High-throughput whole genome sequencing [39–45] or optical mapping technologies [46] have allowed the acquisition of datasets which when analyzed via tailored bioinformatics methods, enable accurate and efficient detection of SVs and CGRs. As such, we have previously reported over one hundred SVs and CGRs in balancer strains of the model organism *C. elegans* [47, 48].

In clinical and biomedical studies, however, SVs and CGRs are still mostly ignored as the interpretation of their involvement in the pathogenic mechanisms is challenging [39]. For single nucleotide variations and indels (small insertions or deletions < 50 bp), we mostly rely on *in silico* tools predicting effect on genes or scoring their pathogenicity, often completed by functional analyses via experimental approaches (model organism or measurement of intermediate phenotype such as RNA or protein levels). However, *in silico* tools to predict the effect of SVs and CGRs are scarce and not yet mature enough to be fully reliable.

For decades, like most Canadian post-secondary institutions, the University of Calgary (UCalgary) has innovated and improved education methods. In 2020, the College of Discovery, Creativity, and Innovation (CDCI) of the Taylor Institute for Teaching and Learning (TI) at UCalgary launched the Undergraduate Research Initiative (URI) to facilitate access to research experiences for undergraduate students and provide support to Instructors to develop and implement innovative teaching methods such as CURE. The Bachelor of Health Sciences (BHSc) program at the Cumming School of Medicine (UCalgary) is an inquiry-based research-intensive undergraduate degree program. We present here a CURE developed for the BHSc as an introduction to Bioinformatics, with the support of the TI, and implemented three times since Winter 2023, for over a hundred students. The research project led by the students consists in the analysis of original and authentic transcriptomic data to explore the functional impact of known SVs and CGRs found in the genomes of *C. elegans* balancer strains.

## Materials and Methods

### Overview

As part of the Bachelor of Health Sciences at the Cumming School of Medicine at the UCalgary, MDSC 301 is the introductory course to Bioinformatics that serves originally second-year undergraduate students of the Bioinformatics program. It was developed as an applied bioinformatics CURE in 2022 and implemented three times. MDSC 301 has accommodated up to 44 students per semester, totaling so far 109 students (Winter 2023 with 30 students, Winter 2024 with 35 students, and Winter 2025 with 44 students). Prior to enrolling in MDSC 301, students are required to complete 6 units in Computer Science at the 300 level or Medical Science 341 or 6 units in Biological Sciences at the 300 level. The CURE is implemented by a single instructor and one to three graduate assistants (a half Teaching Assistant, a Research Coach, and a Graduate Research Assistant). The CURE unfolds over 12 weeks ( January-April) and meets twice a week for 75 minutes. Classes alternate between short lectures and working sessions during which students work in groups and benefit from TAs and instructor mentorship. The course is equipped with a computer science lab, portable wet-bench experimental material, web-based resources including publicly accessible servers (Galaxy), and UCalgary local high-performance computing systems dedicated to students. It is also supported by a molecular genetics research group (Dr. Tarailo-Graovac) and the TI. The main teaching material is in the form of a Lab Book, inspired by previously published CUREs [49]. This document is made available to the students on the first day of the semester and contains all the necessary information related to the CURE: templates, background information, CURE rules, guidelines, protocols, schedule and scripts. The Lab Book 2024 can be found https://github.com/MTG-Lab/MDSC301.

### CURE implementation

MDSC 301 at UCalgary is scheduled once a year in the Winter semester (Figure 1). All materials are prepared in Fall, prior to the class. This includes an update of the Lab Book and acquisition of new transcriptomic datasets. TAs are also recruited in Fall. From January to February, students are guided through the re-analysis of genomics data as in Maroilley et al. (2021, 2023) to acquire the necessary background knowledge in molecular biology, bioinformatics techniques, and research methods. Then students analyze their original transcriptomic data in March and present their findings in a conference organized on the last day of the semester in mid-April. The survey was accessible to students who consented to be part of the study throughout May.

**Figure 1:**
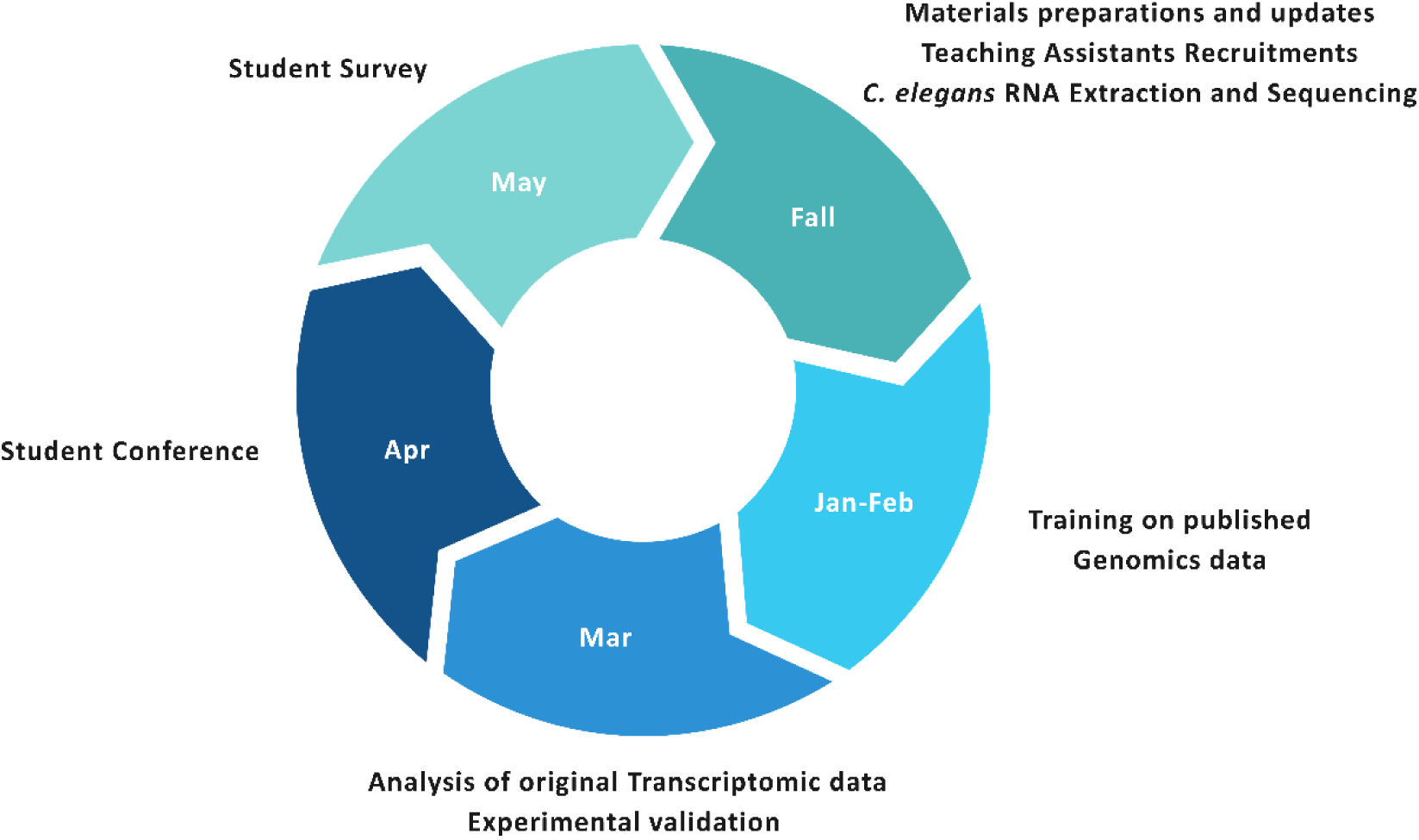
Yearly planification.

### Schedule

The MDSC 301 CURE is implemented over 12 weeks (Figure 2). The first 6 weeks, the students work in group on a mock research project: “How to detect SV in short-read whole genome sequencing data?”. The students repeat analyses done on a strain of their choice published in either Maroilley et al., 2021 or 2022, from which genome sequencing data are publicly available. They are guided throughout the pre-processing of the sequencing data, then apply multiple SV callers, and then assess and compare their accuracy.

**Figure 2:**
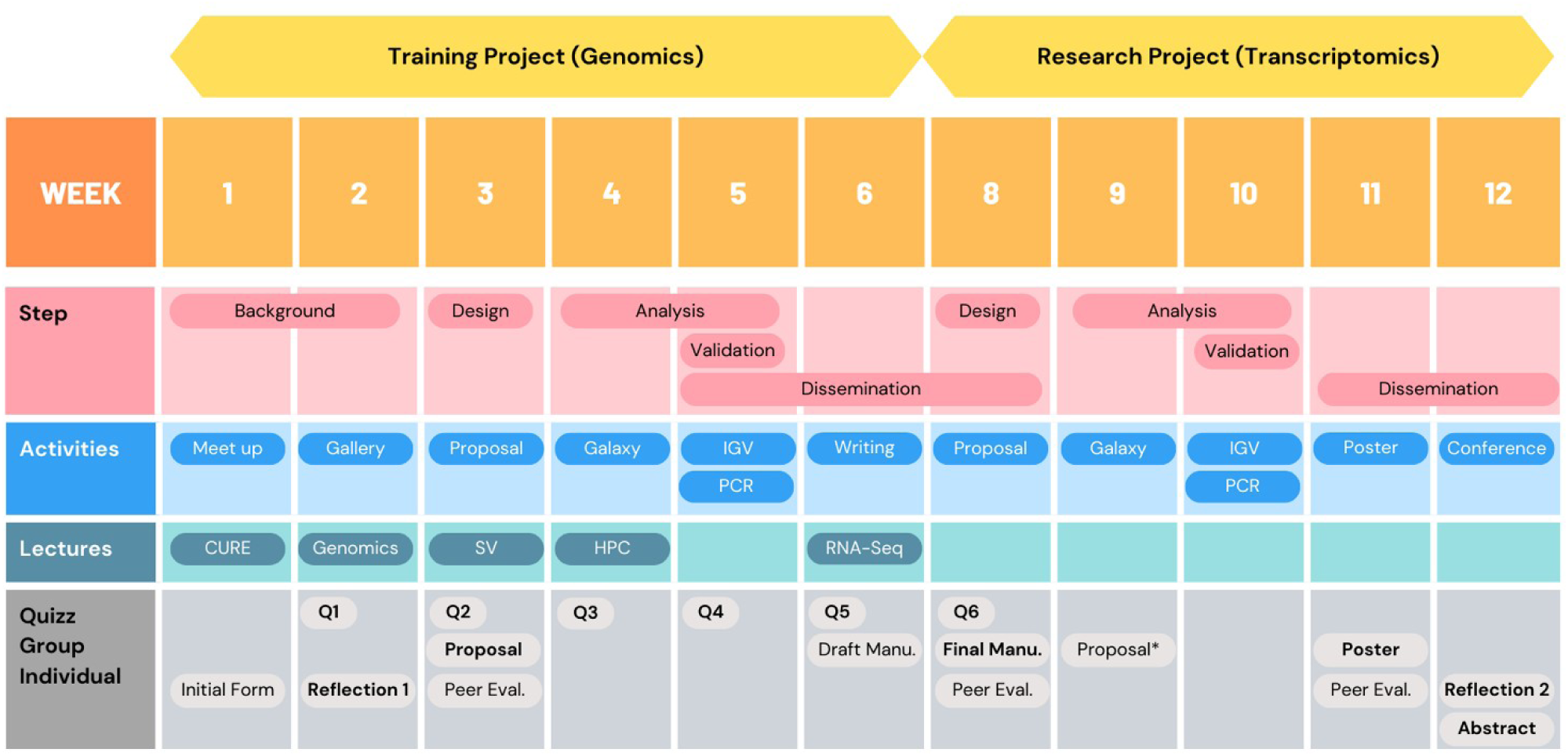
Semester schedule.

In the last five weeks of the semester, the students have access to the original transcriptome sequencing data, from the same strain they used in the genomics project. In that way, they will study the effects of SVs they detected.

### Materials

The main resource for the students is the Lab Book, made available at the beginning of the semester. It was written by the Instructor and the TAs and inspired by previous CURE published online [49]. It contains:

- A presentation of the format of the course as a CURE;

- An introduction to the projects (training and research);

- A set of rules and advice to ensure the success of the CURE;

- A detailed schedule, week by week, with every resource necessary to accomplish required activities and assignments.

- Background summaries: To complete the short lectures and active learning activities, several background documents are available in the Lab Book. They summarize introductory information on the main scientific topics necessary for the students to accomplish their projects: *C. elegans* as a model organism, Genome sequencing, Structural variants;

- Templates: for each research-inspired group written assignment, detailed templates are available. They describe the expected content of each assignment. For instance, for the manuscript due on Week 8, the template summarizes the main sections of a manuscript (Introduction, Methods, Results, Discussion, and References). Each section is explained and the expected content to obtain the full mark is cataloged. As for the Results section, the template explains that only outcomes for the analysis must be reported in this section, and the expected tables and figures are described;

- Project guidelines: For the Training Project (Genomics), the students are guided at every step via a series of protocols with screenshots, command lines and scripts made available. For the Research project (Transcriptomics), the Lab Book offers to the students’ advice, ideas, and explanations, but the students have the freedom of designing their workflow;

- Rubrics for all assignments.

The development of the original Lab Book took about six weeks by the main instructor, helped by some TAs who wrote one or multiple templates or background summaries. Each iteration, the Lab Book is revised and updated based on student feedback and observations made by the Instructor and the TAs while implementing the course.

### Learning Objectives

The course Learning Objectives are multi-faceted as described in Table 1.

**Table 1:**
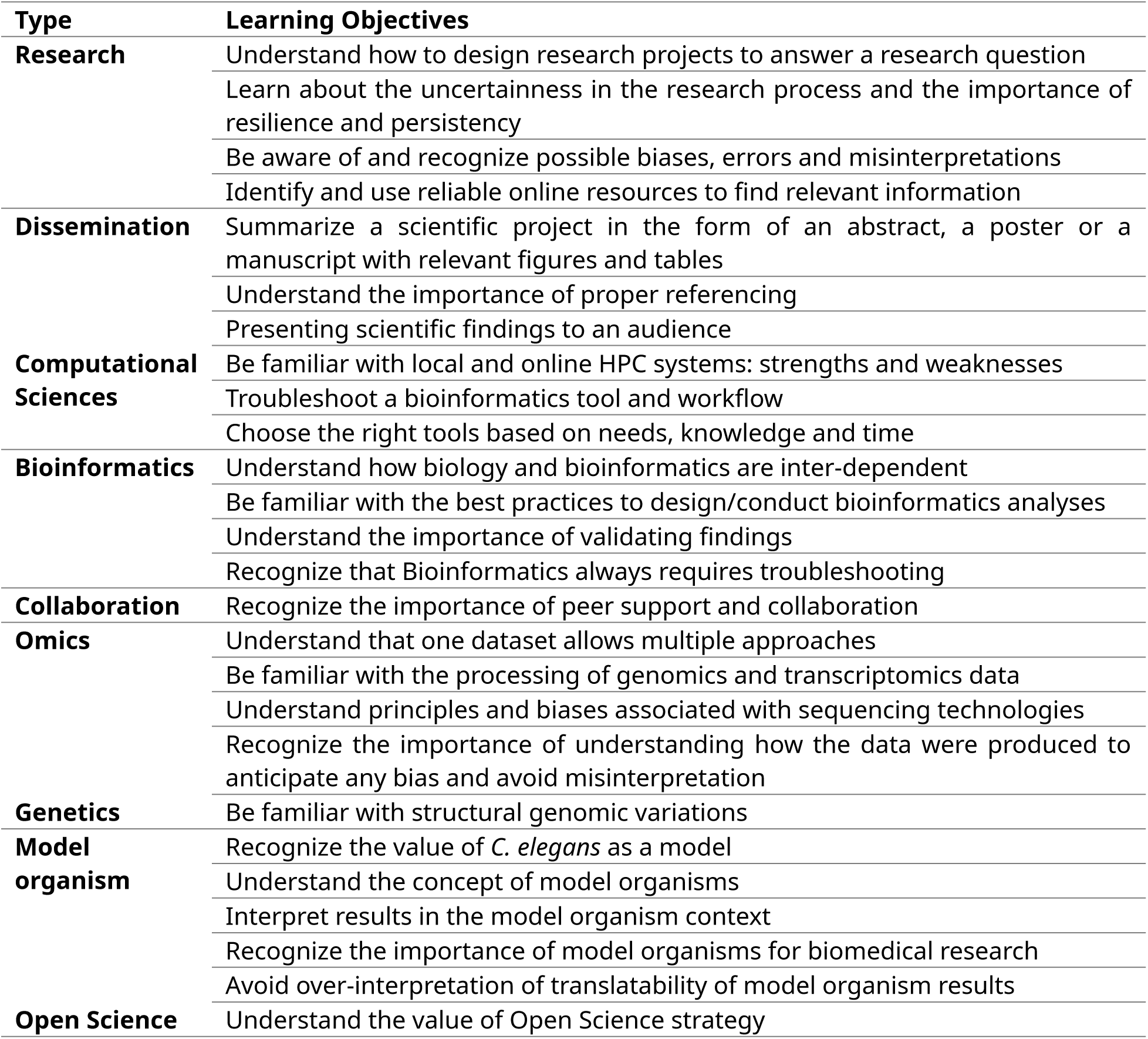
Learning Objectives in MDSC 301.

### Survey

Students enrolled in MDSC 301 in Winter 2023 and 2024 were eligible to participate in the following research study: “College of Discovery, Creativity, and Innovation: Undergraduate Research initiative – What attributes and activities contribute to quality undergraduate research at UCalgary in the course-based undergraduate research programming (CURE), Global Challenges course, Ready for Research, and the Program for Undergraduate Research Experience (PURE)” (Ethics REB20-2110). Participation was anonymous, voluntary, did not require any additional coursework or research duties, and was not associated with performance, assessment, grades, or merit. The student cohorts from 2023 and 2024 were invited to participate in the study and their answers collected from February 2024 to April 2024, and May 2024 to July 2024, respectively.

### Sample collection and data acquisition

At each cycle of the course, three different *C. elegans* balancer strains were analyzed (Table 2)

**Table 2:** Bioinformatics core competencies.

| Bioinformatics Core Competency | Definition | Details |
| --- | --- | --- |
| S1 (Role) | Understand the role of computation and data mining in hypothesis-driven processes within the life sciences | Designing a bioinformatics analysis (pipeline) to answer a biological question |
| S2 (Concepts) | Understand computational concepts used in bioinformatics | Pipeline design, tools, file formats (Fastq, Fasta, Bam, VCF) |
| S5 (Tools genomic) | Be able to use bioinformatics tools to analyze genomic data | FastQC, Trimmomatic, MarkDuplicates, BWA, SV callers, IGV |
| S7 (Tools expression) | Be able to use bioinformatics tools to analyze gene expression data | Cutadapt, STAR, FeatureCounts, HTSeq-count, Limma, DESeq2, OUTRIDER, FRASER |
| S13 (Scripting) | Know how to write short computer programs as part of the scientific discovery process | (OPTIONAL) Bash, R, python |
| S14 (Software) | Be able to use software packages to manipulate and analyze bioinformatics data | See S5 and S7, Spreadsheets |
| S15 (Computational environment) | Operate in a variety of computational environments to manipulate and analyze bioinformatics data | Web-based, (OPTIONAL) Unix/Linux command Line |

For the Training project, genomics data were collected from Maroilley et al., 2021 and 2023. For iteration of the CURE, N2 and Hawaiian (CB4856) strains were used as a control, and three balancer strains were made available to the students:

- 2023: BC986, BC4586, and VC109
- 2024: CZ1072, MT690, and SS746
- 2025: BC1217, GE2722, and RW6002

For the Research project, total RNA was extracted from a pellet of about 1,000 worms with a Trizol-chloroform based RNA extraction protocol followed by in-column DNAse treatment ZYMO Research RNA Clean & Concentrator kit. (ID: R1013). Purified RNA was eluted in DNAse-RNAse free water provided by the manufacturer (ZYMO Research). RNAs were depleted from ribosomal RNAs. mRNAs were then sequenced at the Centre for Applied Genomic Next Generation Sequencing Facility (SickKids, Toronto, Canada) with Illumina NovaSeq 6000. For each student cohort, N2 was used as a control.

### Genomic data analysis

The genomic data are 150-bp Illumina short reads derived for the genome of *C. elegans* balancers stored in fastq files. Raw data were preprocessed as in Maroilley et al., 2021 and 2023, on the online HPC system Galaxy (Galaxy Europe at https://usegalaxy.eu). Each student created a free account on Galaxy Europe (https://usegalaxy.eu/). Data were uploaded in “History” by the instructor and later shared with the students. The quality control of the data was checked with FastQC [50]. Trimmomatic [51] was then ran on the raw data to trim reads from bad quality bases, discard unpaired reads, and remove adapters. Duplicated reads were marked with MarkDuplicates (Picard) [52]. Reads were then aligned on the *C. elegans* reference genome (ce11) using BWA-MEM [53]. Quality of the alignment was evaluated using FastQC [50]. Structural variants (SVs) were called using Delly [54], Lumpy [55] and Basil [56], available on Galaxy. SVs were also called with Manta [57], SeekSV [58], GRIDSS [59], and TIDDIT [60], ran on a local HPC system (TALC, University of Calgary). Calls were filtered based on quality scores of each caller, overlapping between callers, and visual inspection of breakpoints using Integrative Genomic Viewer (IGV) [61].

### Transcriptomic data analysis

The transcriptomic data are 150-bp Illumina short reads derived for the genome of *C. elegans* balancers stored in fastq files. Transcriptomic data pre-processing was implemented on Galaxy Europe. Quality of the raw data was assessed with FastQC [50]. Reads were trimmed with Trimmomatic [51]. Adapters er removed with either Trimmomatic [51] or Cutadapt [62]. Reads were aligned to the reference genome with STAR [63] or HISAT2 [64]. Manual analysis was performed on IGV [61]. Read counts were obtained using featureCounts [65] or HTSeq-count [66]. Differential expression analysis was made using limma [67], allowing the absence of replicate in the balancer group, or DESeq2 [68], either on Galaxy Europe (https://usegalaxy.eu/) or R (https://www.r-project.org/).

### Experimental Validation

Breakpoints were validated by PCR on DNA. Gene expression levels were studied via RT-PCR on cDNA. The PCR reagents were mixed as follows: 15 µL or water, 4 µL of 5X EZ PCR Master Mix, 0.4 µL of 10 µM primers, and 0.6 µL of worm lysate. The PCR thermocycler was run as follows: initial denaturation 95°C for 3 minutes, denaturation 95°C for 30 seconds, primer annealing 54°C for 30 seconds, extension 72°C for 45 seconds, and final extension 72°C for 5 minutes. Denaturation-annealing-extension was repeated 35 times. Gel electrophoresis performed for 30 minutes at 100V using a 2% agarose gel loaded with 10 µL of PCR mix (1:5 dye:sample).

## Results

### The MDSC 301 CURE has all characteristics of a CURE

CUREs are defined by five main characteristics that our course fulfilled [23]:

1. Scientific process: The students are going through the scientific process twice over the course of this CURE, once during the Training project (Genomics), and once during the Research project (Transcriptomics) with more guidance the first time. Each time, students have access to a short list of research and review papers in which they can find relevant information to design a project ensuring a specific question (“How to detect SV in genome sequencing data?” and “Which effect can SV have on gene expression?”).
2. Discovery: The data used during the Research project (Transcriptomics) are original, obtained by a research laboratory and not yet analyzed. Any result obtained by the students is in this way an authentic discovery.
3. Relevance: SVs and CGRs are known to change the structure of large genomic regions, and it is to be expected that this would have consequences on downstream phenotypes (RNA, protein, and visible phenotypes). However, there are currently no standard ways to interpret the impact of an SV computationally and SVs are often ignored in genetics and genomics studies. By exploring the effect of many various SVs, we can pave the way to developing prediction tools.
4. Collaboration: Research projects have been designed in a way so that goals can be achieved via the collaboration of at least four students. In addition, we encourage collaboration between teams.
5. Iteration: During the Training project (Genomics), students are following a protocol provided in the Lab Book. By doing it themselves, students face the day-to-day challenges in Bioinformatics and get to re-try and troubleshoot (learning by failure and retrying). For the Research project (Transcriptomics), students design their analysis, involving making choices and having to adjust them (revising thinking).

### A design addressing core competencies and translational skills

The Network for Integrating Bioinformatics into Life Sciences Education (NIBLSE) has published a list of 15 bioinformatics core competencies [69]. Table 2 summarizes the seven Bioinformatics core competencies students are familiar with MDSC 301 CURE. S13 and part of S15 (Table 2) are based on activities in the Training Project (Genomics) made optional for students with previous bioinformatics or computational sciences experience, or particular interest in the usage of command-line, scripts, and UNIX systems (see “The challenge of introducing applied bioinformatics to students with different university backgrounds” section for more details).

As students experience research, they develop or reinforce interdisciplinary and transferable skills. Table 3 summarizes the transferable skills supported by the MDSC 301 CURE.

**Table 3:** Transferable skills.

| Skills | Definition |
| --- | --- |
| Collaboration | Work respectfully and efficiently with others, plan a collaborative project with milestones. |
| Communication | Learn and share information by presenting orally and in writing, listening, and interacting with others. |
| Creativity | Elaborate on new approaches. |
| Critical thinking | Actively and skillfully conceptualize a solution and apply a method. Analyze, synthesize, and evaluate results/information to draw a reasonable conclusion. |
| Digital skills | Use digital technologies (computers, high performance computing system, online resources). |
| Literacy | Find, understand, and use information presented through words. |
| Numeracy | Use mathematical information. |
| Problem Solving | Identify an issue, know how to look for (online, literature, peer support), find, implement and assess a mitigation. |
| Mentoring | Peer-mentoring based on individual strengths and university background. |

### The challenge of introducing applied bioinformatics to students with different university backgrounds

Originally part of the Bioinformatics program in the Bachelor of Heath Sciences at the Cumming School of Medicine as a second-year mandatory class, the MDSC 301 is opened to students from other programs and other years. On average, over the three years of implementation, only a third of the students were second-year Bioinformatics programs students. Other students were third- or fourth-, fifth-year students from Biological Sciences, Cell and Molecular Biology, Computer Science, or Biomedical Sciences. Their experiences in Bioinformatics were either none, a block-week introductory class, a few classes, or summer internships in research laboratories working on projects involving Bioinformatics.

To address the challenge of conducting a research project with Bioinformatics approach that would be accessible and engaging for all students, several measures were implemented:

1. At the beginning of the semester, students were asked to complete a survey answering questions to evaluate their experience in Bioinformatics, Genetics, and research (See Supplementary Information). This allows the instructor to adjust the curriculum to the current cohort of students.
2. Provide a Lab Book with tutorials for every step of the Training Project (Genomics) and guidelines for the Research Projects (Transcriptomics) (https://github.com/MTG-Lab/MDSC301).
3. The core of the Training Project (Genomics) was adapted to the free online platform Galaxy (Europe), allowing anyone with no experience in UNIX systems, command-line or coding to run Bioinformatics analyses. The most popular tools have been made adapted on Galaxy for easy utilization.
4. As an option for students majoring or minoring in Bioinformatics, or with experience or interest in using command-line, scripts and UNIX system, we provide access to a local High-Performance Computing system (TALC, University of Calgary). As such, when other students run SV callers on Galaxy, volunteers would run similar callers pre-installed on the cluster. A SLURM script template is provided to facilitate running a job and guidance during class hours. The instructor and TAs are available during office hours and via email to students. To avoid students feeling pressured in choosing one of the options, grades were not affected by the system used.
5. Teams were built based on answers received via the initial survey (see Supplementary Information). In brief, we collect names for students they would trust to work with, but also their experience in research, Bioinformatics and Genetics, and their self-assessed skills in writing and presenting. As such, we built teams of four students as follows: i) accommodating group request; ii) one member of each group should have prior Bioinformatics/research experience; iii) one member of each group should have good writing skills.
6. Hiring teaching assistants with Bioinformatics background to ensure mentorship to students who need it.

### Fostering (smooth) collaboration

Bioinformatics suffers from stereotypes often leaving students thinking that Bioinformaticians or researchers in Bioinformatics work alone. However, collaboration is essential in Bioinformatics, as it is in research, but even more as the field is rapidly evolving and knowledge is not always recorded in books or published as articles. In addition, collaboration is a transferable skill, valuable in any path students will be joining after their degree.

To foster as much collaboration as possible in the class, most of the research-like assignments were to be done in groups. MDSC 301 implemented then a Team Based Learning approach: the groups are formed by the instructor and team members evaluate each other’s contribution. This approach was announced to the students in the first class to give them time to raise any concern and get familiar with the concept before any group assignments. Additional information, advice, and good practices were available to the students in writing in the Lab Book. In brief, the main research-assignments were to be done in group: the Training Project (Genomics) proposal (10% of the grade), the Training Project (Genomics) manuscript (20% of the grade), and the Research Project (Transcriptomics) poster (20%).

To give the groups the opportunity to get to know each other before group assignments, we formed the teams as early as possible in the semester (second week), and we planned active learning activities in groups to get team members to work together without the pressure of group assignments for a few weeks. In addition, group work can only be successful with efficient organization including task allocation and milestones. But such planification requires understanding what precisely the assignment unfolds. We have then decided to guide the students in their project management of the Training Project (Genomics) to avoid setbacks and pitfalls that students would encounter just due to bad planning. As such, we give them an example of how to divide the tasks in their group to allow each student to work in parallel on similar but not overlapping tasks. This project management allows mutual assistance in groups but also among groups but still gives each student responsibility for an essential part of the project.

We used individual feedback to give an opportunity to each student to report any issue in their group and ask for help individually. Feedback was collected at multiple time points during the semester through different formats:

1. Peer-evaluation: With each group assignment due, students were due for an individual peer-evaluation. They would assess the engagement, reliability, and efficiency of each of their teammates and themselves. The peer-evaluation would affect a student grade only if on average a student would receive a peer-evaluation lower than 10/20. Otherwise, it was used to spot any minor issue in the group that would have arisen.
2. Reports: Reports are individuals and questions are deliberately oriented to make students reflect on their performance as an individual and as a teammate. Some questions are also designed to gather information about group organization while giving advice to the students about how to improve group work (see supplementary information).
3. Anonymous feedback: As not every student would feel comfortable speaking up, we also made available throughout the semester a link to an online survey that allowed anonymous feedback.
4. Daily availability: the instructor and the teaching assistants were always attending classes, offering office hours, and responding in a timely manner to emails to create and maintain trust for students that they feel supported and welcome to reach out.

### Engaging students in an active learning journey

To acquire the necessary knowledge to conduct the Research Project (Transcriptomics), we implemented various learning activities to engage the students in an active learning process, but also to satisfy all learning styles.

First, all lectures were less than 30 minutes, aiming to deliver the main information necessary to the execution of a part of the project. Slides are made available to the students and are the main resources for quizzes. Then, summaries of basic knowledge were included in the Lab Book, giving the students the opportunity to always get back to this resource. If students wanted to come prepared in class or to explore more deeply a subject, for each session, optional readings under various formats were listed in the Lab Book (videos, free textbooks, free research or review articles). Finally, in any literature search that was necessary, a short list of meaningful articles was always proposed to offer to guide the students in their search while providing foundation information.

Several active learning activities were implemented:

1. “Meet up with the worms and the team”: To introduce the students to the concept of research with model organism (here *C. elegans*, a 1mm-long roundworm), we organize a visit of a research group in the classroom. For one hour, students observed worms under the microscope and discussed with graduate students and postdocs who work with *C. elegans* daily. In addition, we invited graduate students and postdocs working essentially on computational Biology techniques. The meet-up was guided via an individual written assignment of six questions, designed to make students ask relevant questions to the scientists regarding research, Bioinformatics and *C. elegans* (see supplementary information).
2. Walking Gallery: For students to acquire the necessary scientific knowledge to conduct a genomics analysis of SVs, we implemented a walking gallery activity, where students are building the content of the session. Prior to the session, students were asked to read pre-selected articles (four per group, so one per student) in the light of the themes they would have to discuss during the activity (themes available in the Lab Book). In class, students will go in groups from one theme to another (written on posters or white board) to record in writing any piece of information relevant to the subject they would have read in their paper. Each group who would follow would build upon what was already written. This activity was implemented twice: once for genomics, once for transcriptomics. As such, students were even more prepared the second time and the session was even more prolific.
3. As part of the research projects, the students tried to validate some of their findings experimentally. We acquired the miniPCR Bio kit, the miniPCR thermocycler and the blueGel electrophoresis system, a portable technology allowing to perform PCR in any environment, including a classroom (https://www.minipcr.com/). To fit in with the 1.15-hour session, students were instructed to design primers; however, the primers were designed and ordered by the instructor ahead of time. DNA or RNA extraction, PCR mix and gel preparation were performed by the graduate assistants before the class. Students loaded the gels with their samples. Gel was run in the classroom in front of the students, who, thanks to the blueGel electrophoresis system, could follow in real time the progression of the experiment. An example of gel is available in Supplementary Information.
4. Mini-symposium: Toward the end of the semester, students are engaged in a demanding research project as they analyzed the transcriptomics data to study the effect of SVs on gene expression. To allow them to fully focus on their research, we do not impose a final exam. In term of final assignment instead, we require the design of a poster. To promote engagement and genuine motivation, we organize a one-hour mini-symposium like Summer Research symposiums on the last session of the course. The event is advertised to gather students and researchers at the event and give a chance to the students to present their work and findings to a varied audience. The presentation of the poster during the event is not subject to any evaluation for the students’ participation, as it was a professional conference.

### Supporting students in research-like assignments

To allow students to experience the research process from designing a project to disseminating results and conclusions, we planned research-like assignments: proposal, manuscript, abstract, and poster. However, such assignments can be overwhelming for undergraduate students as each format has a set of rules and best practices that scientists usually learn-by-doing over several years in a lab. In the context of a CURE, it was also important to make sure that the assignments would not take over the bioinformatics analysis that students were conducting. Then, to allow easy time management for students, and guide them into the production of research-like assignments while learning the best practices, we created detailed templates and guidelines made available in the Lab Book (see Lab Book).

In addition, we planned a major part of each session in class to be a time for students to work on their project and assignment. As such, any question or issue arising can be addressed immediately by the instructor, the teaching assistants or peers, avoiding any delay in the completion of the assignments. During work sessions in the class, the teaching time will also go around from one team to another to ask for updates and offer help. We found this approach important to reach students reluctant to ask questions. It was also critical in noticing teams that were encountering delays and providing personalized support to help them to be back on track. Work sessions also favored collaboration inter-teams, even more encouraged by the instructors when some teams would struggle obtaining a file for technical reasons, “colleagues” working on the same sample were asked to share their file allowing everyone to pursue their analysis.

For each research-like assignment, early deadlines were set to allow students to pre-submit a draft of their assignment to receive feedback before finalization and final submission. The feedback would highlight any missing requirement, advice on the format and the content, review the quality of the writing for the available sections, and suggest techniques to improve/complete the draft and any remaining analysis in a timely manner.

Finally, the rubrics were made available and reflected the marking system. We graded the research-like assignment based on their alignment with the requirements described in the template – “was all required information presented in the appropriate format?” The grade was not affected by the scientific findings, as an attempt to allow students to conduct their analysis without the pressure of finding the “right answer” as in usual assignments. Here, we favored the intellectual process behind the analyses and the dissemination of the results over the accuracy of the results. We also encouraged and guided the students in explaining, justifying, and reflecting on possible biases and limitations in their chosen methods as is done in scientific publications.

### Helping students evaluate their learning

As previously described, the MDSC 301 CURE allows students to acquire, develop, or improve bioinformatics core competencies, and transferable skills. As a CURE can be confusing in terms of how it will beneficiate students who do not wish to pursue research and/or in the specific field of research in which the project fits, we implemented resources similar to metacognitive assessments to favor students’ self-assessment at the beginning and the end of the semester, encouraging them in reflecting on their journey and their learnings.

First, the Initial Survey (see Supplementary Information) asked several questions to each student regarding their outlook of Bioinformatics and research, current questions, and expectations for the course. Then following the “Meet up with the worms and the team” activity, previously described, students are required to complete their first individual assignment (Reflection 1, see Supplementary Information). The format is a series of questions regarding the observation of worms and the discussions with model organism biologists and computational biologists. The questions were encouraging self-reflection regarding how this experience changed, or did not change, their understanding of research, Bioinformatics, and model organisms. The final individual assignment, called Reflection Assignment 2 (see Supplementary Information) was designed to mirror the first two self-reflection activities by asking similar questions to the students at the end of the semester and allow them to acknowledge their growth in terms of scientific knowledge, transferable skills, but also as a person.

### Impact of MDCS 301 CURE on student experiences

As part of the study REB20-2110, a survey was conducted for voluntary students’ post-course (see Methods). Overall, nine (4 students in 2023, 5 in 2024) participated (13%). In sum, students who participated in the survey reported that the MDSC 301 CURE had positive effects on their learning. Students also felt that the CURE had a positive effect on their engagement in their program and clarified their interests for future studies or career (Figure 3, Table 4).

**Figure 3:**
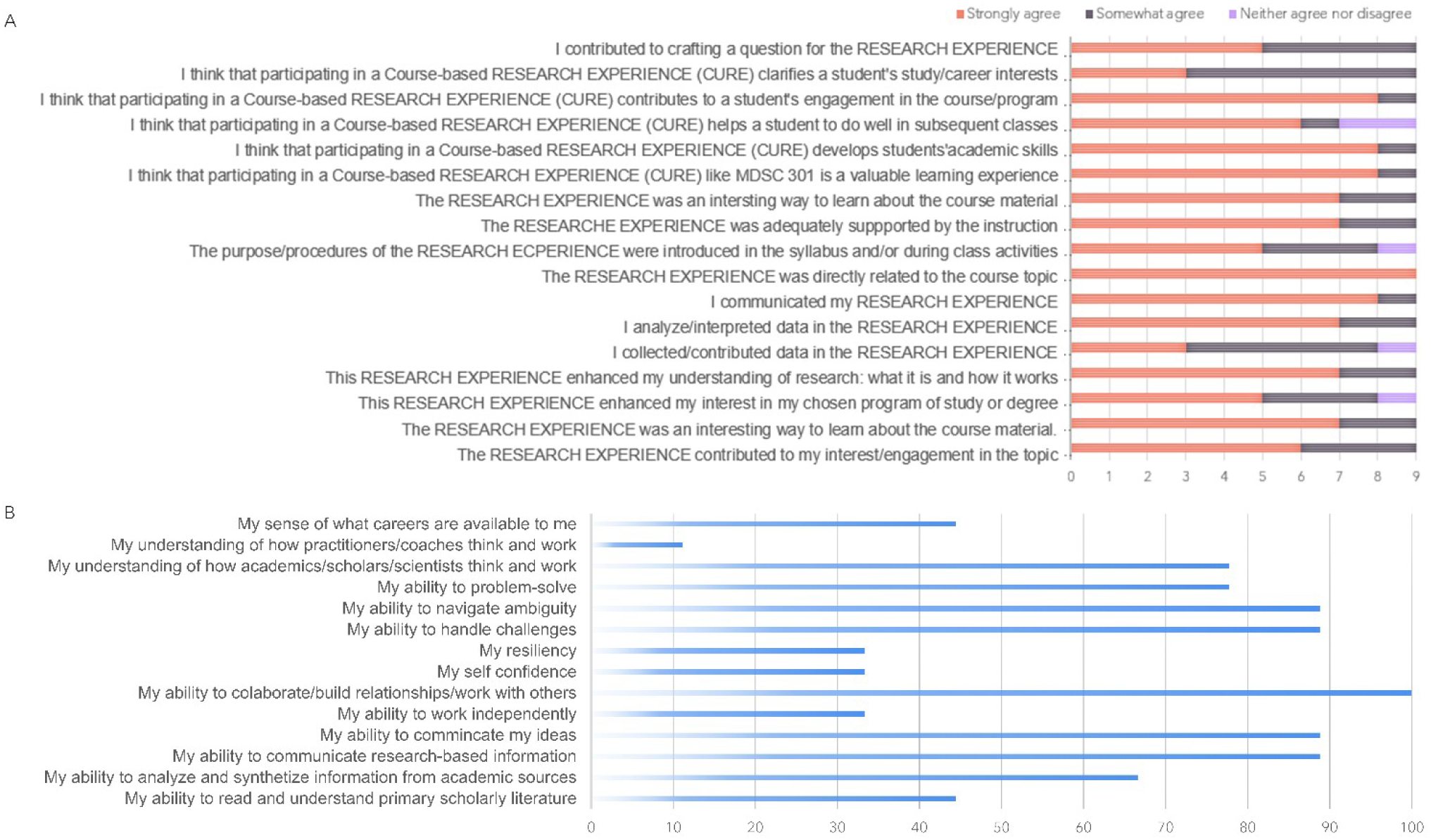
Survey results.

**Table 4:** Open statements.

| Open statement | Sample responses |
| --- | --- |
| **As a result of this RESEARCH EXPERIENCE, I...** | Learned more about Bioinformatics and what encompasses as a field. |
|  | Have a better understanding of the research process and how research is conducted in the real world. |
|  | Have much more thorough understanding of research approaches in Bioinformatics, specifically in RNA sequencing. |
|  | Acquired a deeper understanding of the importance of team collaboration in scientific research, fostering teamwork, negotiating, and compromise while working towards a common goal of producing high quality research output. |
| **Participating in a RESEARCH EXPERIENCE meant...** | I was able to work with my peers and collaborate to produce a final project. |
|  | That I got to actually apply what had been taught and not simply absorb theoretical information. |
|  | I got an opportunity to work in a team to complete rigorous research projects in an area of personal interest. |
|  | Developing essential skills such as critical thinking, problem-solving, and effective communication within a collaborative team setting, fostering a holistic approach to learning beyond traditional coursework. |
| **This RESEARCH EXPERIENCE helped develop...** | Skills in creating a pipeline and analysis related to it. |
|  | Skills such as making a poster to present relevant findings. |
|  | My passion for Bioinformatics and for research as a whole. |
|  | My ability to perform RNA sequencing analyses. |
|  | My presentation skills significantly, particularly through the creation and delivery of our poster presentation during the mini-symposium. It honed my ability to distill complex information into a concise and visually engaging format, as well as to effectively communicate our findings for a diverse audience. It offered a safe place to refine my skills in a setting that mirrored professional conferences, ensuring I entered future endeavors equipped with confidence and proficiency in scientific communication. |

### An example of scientific discovery: *eT1* disrupts more than the expected *unc-36*

*eT1(III;V)* is a reciprocal translocation used to balance the right end of chromosome III and the left end of chromosome V in *C. elegans*. Breakpoints are III: 8,200,764 and V: 8,930,675. *eT1(III;V)* is easily traceable in a colony as its presence produces an uncoordinated phenotype. It was suspected that this phenotype was due to the disruption of the *unc-36* gene, situated at the chromosome V breakpoint. However, little was known regarding the potential effect of the chromosome III breakpoint, as it is located in an intergenic region.

We sequenced the transcriptome (bulk RNA) of the BC986 strain, carrying the *eT1(III;V)* reciprocal translocation. Visual observation of the RNA-seq alignment in the genome viewer Integrative Genomic Viewer (IGV) performed by the students helped further understand the effect of *eT1*.

At the chromosome III breakpoint, the transcriptome data show a different expression of the *unc-36* gene, when compared to control: in BC986 only exons 1 and 2 are well covered by reads, while the rest of the *unc-36* presents almost no coverage. In comparison, in N2 (control), the level of coverage is stable throughout the whole transcript. This suggests a truncation of the transcript, which aligned with the location of the breakpoint, just left to exon 2 (Figure 4).

**Figure 4:**
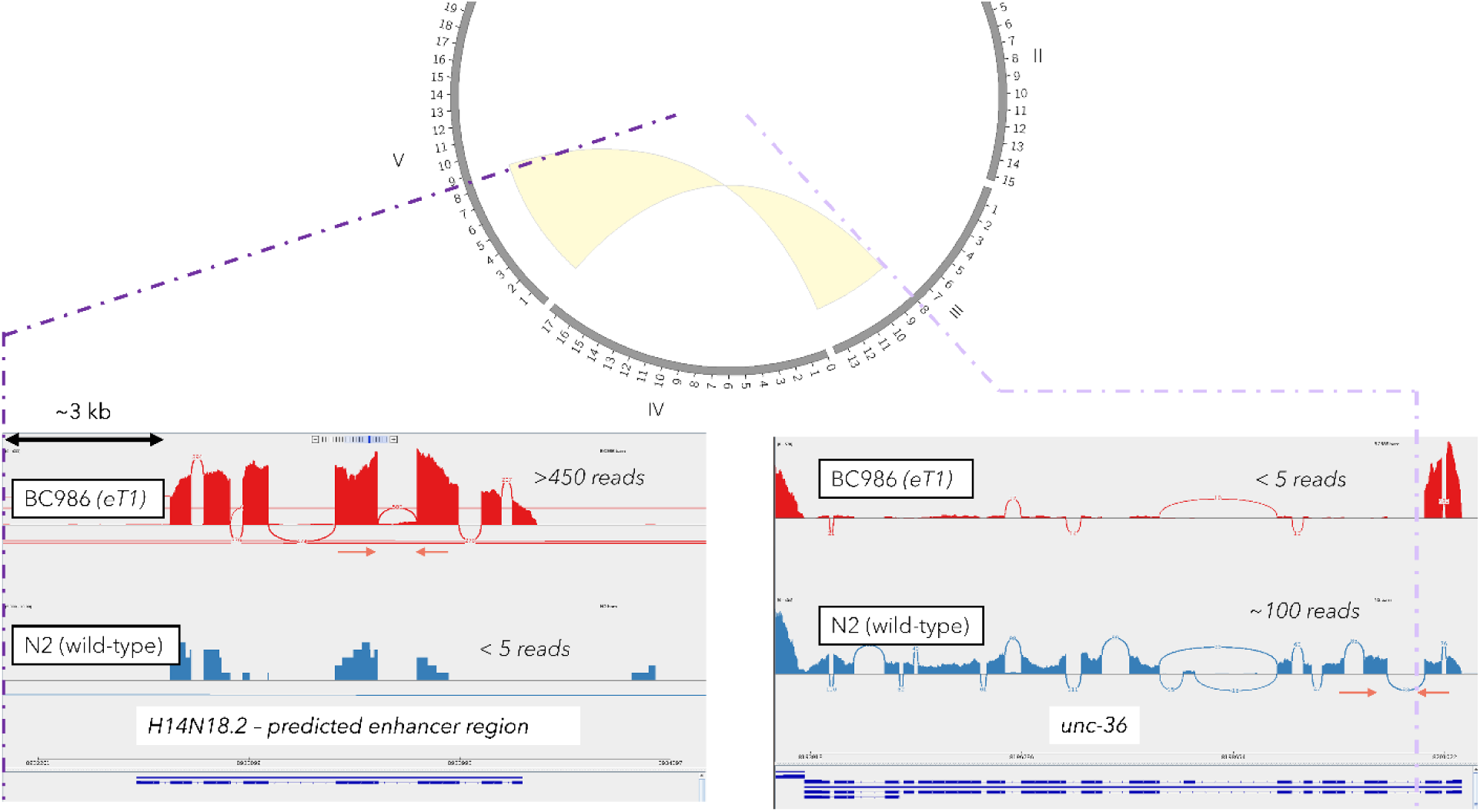
eT1 on BC986 genome, and effect on its transcriptome at both breakpoints.

The chromosome V breakpoint is not overlapping any known annotated genetic element (gene or non-coding). However, a visual analysis of the RNA-seq alignment shows a difference notable variation in the coverage of *H14N18.2*, predicted enhancer region, located at about 3 kb from the *eT1* breakpoint (Figure 4, left panel). The pattern of coverage shows clearly distinctive exons that are barely covered on the N2 control. This suggests that the *eT1* breakpoint induces an expression of *H14N18.2*, usually not expressed in N2. As *H14N18.2* is understudied, its expression in an *eT1* balancer-containing strain gives material for further investigation of this genetic element in the *C. elegans* genome.

## Discussion

The fields of Bioinformatics, Computational Biology, and Data Science are exponentially expanding with the advent of high-throughput technologies and Artificial Intelligence. Multi-disciplinary training is necessary, but the offer remains limited and often implemented in post-secondary institutions for graduate students. This impacts directly the limited interest of students in any field related to Bioinformatics, which suffers from a stereotype of a narrow, impersonal, and unsuited field for those who wish to work on a human level, discouraging students from pursuing these paths. Implementing additional introductory courses to Bioinformatics and programs at undergraduate levels would foster interest of students, as they would have a better understanding of what Bioinformatics involved. But it has been shown that in the forms of lectures with theorical content, introductory Computer Science courses play a big role in discouraging students from majoring in computer science, especially women. To foster student engagement in Bioinformatics-related career, it is then of the utmost importance to improve introductory curriculum, tailoring how we teach Bioinformatics by “marketing” the courses [69]. Here, we design and implement a CURE (Course-based Undergraduate Research Experience) as an introduction to Bioinformatics, in the form of a research project with applied Bioinformatics techniques on omics data. In brief, we combined multiple approaches to ensure engage all learning-type students: authentic research project, original data, active learning activities, research-like assignments and, open-to-public mini-symposium. While conducting a post-course survey, we showed that students’ interest and understanding in the field have improved.

Undergraduate Computer Science programs remain remarkably un-diverse, with women only earning 18% of computer science bachelor’s degrees in the United States for instance. Women and other minority groups are also dismal in the computer science professions, with fewer female authors, in sex or gender, on research papers, as compared to biology in general (∼20-30%) [70] and a persistent under-representation of minorities in the field [71, 72]. In our CURE, we aim to support students from underrepresented groups by creating an inclusive learning environment, where all students feel that their differences are valued and respected, have equitable access to learning and other educational opportunities, and are supported to learn to their full potential. As such, suggested readings were exclusively free resources, students had access to university computers, and the analyses were feasible via an online free platform (Galaxy) or a computing system available to students at UCalgary.

Here, not only are we showing the impact of a CURE in Bioinformatics on students, but we also intend to disseminate our curriculum, and all materials and strategies developed to allow implementation of this CURE in other programs or facilitate the design of other CUREs by providing ideas, support and resources. Indeed, designing a CURE from the start can be intimidating and instructors might be reluctant due to the charge of work it might represent. We have then chosen to share with the community our materials to reduce this burden and favor the emergence of new CUREs. Our CURE was inspired by Villa-Cuesta & Hobbie [49] in its form but developed around an original research project developed in the Dr. Tarailo-Graovac laboratory and based on instructor competencies. By providing all resources, and in the era of Open Science with most datasets being publicly available, even instructors with not the exact skillset for this project could implement this course at least partially.

Any data science research project is designed around three essential elements: a research question, a dataset, a method of analysis. To design a successful CURE as an Introduction to Bioinformatics, the main caveat is to think that students need independence to conduct their research. On the contrary, undergraduate students in a class, just as summer students in a laboratory, need a reliable support system and a framework for successfully conducting their research efficiently. As such, it is important to design a project by providing the students with at least two of the three essentials (question, method, data). In MDSC 301, we choose to have students working on the same research question (effect of SV on gene expression), and to provide them with the datasets. They could then innovate on the methods to analyze the data to answer the research question. One could choose to let the students find their own research question but provide a specific dataset or a repository (e.g., TAGC) and a method (e.g., a differential expression analysis). Or students could be provided with a method to apply a specific question, but they would have to find their own dataset.

Our design was based on research in which the instructor was involved. As such, CURE provides the perfect opportunity to combine both research and teaching activities often required from academics. It becomes then not only beneficial to the students, but also to the researcher beyond teaching. Indeed, research and discovery are made in such a way that it is publishable, a reward for the students and the researcher.

## List of abbreviations

CGR: Complex Genomic Rearrangement
CURE: Course-based Undergraduate Research Experience
SV: Structural Variant
UCalgary: University of Calgary
C. elegans: Caenorhabditis elegans
CDCI: College of Discovery, Creativity, and innovation
TI: Taylor Institute for Teaching and Learning
URI: Undergraduate Research Initiative
BHSc: Bachelor of Health Sciences
TA: Teaching Assistant

## Ethics approval and consent to participate

The post-course survey was approved by the University of Calgary (REB#: REB20-2110) for the study “What attributes and activities contribute to quality undergraduate research at UCalgary in the course-based undergraduate research programming (CURE), Global Challenges course, the Program for Undergraduate Research Experience (PURE), and the Ready for Research microcredential”. Clinical trial number: not applicable.

## Availability of data and materials

Materials from the Winter 2024 MDSC 301 (second iteration) can be found at https://github.com/MTG-Lab/MDSC301. The genomics datasets were included in Maroilley et al., 2021. The RNA-Seq datasets generated and analysed during the current study are available in the GEO repository GSE292388.

## Authors’ contributions

Research design and conceptualization were by T.M. and M.T-G.; CURE design was by T.M. with the help of V.R.A.B and R.M. Data analysis and interpretation were performed by F.A., S.A., A.A., D.A., M.A, S.A., K.A., R.B., B.C., L.D., P.E.N., E.G-V, Z.H., S/H., G.J., N.K., I.L., C.L., T.L., S.M., S.P., S.P., R.Q., F.R., B.S., S.S.A., H.A.T, R.W., C.M.W., S.Z with the help of T.M., V.R.A.B, and R.M. C.D. and S.F. prepare the strains and extracted the RNAs. S.F. set up the experimental validations in class; molecular analysis and confirmation of the variants were by T.M., F.J., V.R.A.B., X.L., and L.O.; and original draft preparation was by T.M., S.F., M.E., D.M., and M.T-G. All authors have read and approved the submitted manuscript.

## Acknowledgements

We want to thank Kyla Flanagan, Rachel Stuart, Kara Loy, and the CURE program implemented at the Taylor Institute for Teaching and Learning at UCalgary for their support in the development and implementation of the CURE via meetings, workshops, and funding for Research Coaches in 2023 and 2024. We want to thank the Teaching assistants and Graduate Assistants who helped with the implementation of the class: Diogo Marques, Shreya Tomar, Jinsu An, and Zeyad Abouyoussef. The authors acknowledge the support of the Freiburg Galaxy Team: Person *X* and Björn Grüning, Bioinformatics, University of Freiburg (Germany), funded by the German Federal Ministry of Education and Research [BMFTR](https://www.bmftr.bund.de/) grant 031 A538A [de.NBI-RBC](https://www.denbi.de/) and the Ministry of Science, Research and the Arts Baden-Württemberg (MWK) within the framework of LIBIS/de.NBI Freiburg.

## Authors’ information

