## Supplementary Information for "A Course-Undergraduate Research Experience (CURE) to explore the effect of structural variants on gene expression in *C. elegans* balancers"

#### Initial survey – Evaluating students experiences

1. What program are you in?
2. What is your degree?
3. In which year are you in?
4. List of the classes you have previously taken in Bioinformatics
5. Briefly explain your contributions in projects with Bioinformatics analyses in a lab or in industry, if any
6. List the classes you have previously taken in Genetics
7. Briefly explain your contributions to projects on model organism, if any
8. Are you comfortable working in groups?
9. Are you comfortable talking in public during presentations?
10. Are you comfortable writing essays in English?
11. Describe yourself as a teammate (you can pick multiple answers)
    a. I am not afraid of taking responsibility and making choices
    b. I usually follow the group consensus
    c. I often disagree with the group and have a hard time compromising
    d. I always fulfill whatever tasks the group has assigned me on time
    e. I usually have a hard time respecting deadlines
    f. I like group work when everyone has an independent task to do
    g. I like group work when we do everything together and discuss everything together
    h. I do not like when a team member take initiative without discussing it with the group
    i. I always try to make sure that my contributions equal others’
    j. I think that work groups always imply that one will work more than the others
    k. I like to be part of discussions and the decision-making process
12. List classmates you would prefer to work with, if any
13. Three things you expect from MDSC 301
14. Two things you already know about Bioinformatics
15. One thing you are wondering about Bioinformatics

### Report

1. What part of the project is your responsibility and what progress did you make?
2. Do you feel that the work has been appropriately distributed in your group?
3. Are you confident/anxious about this group assignment?
4. What communication plan did you and your team put in place?
5. Did you establish deadlines?
6. Did you establish regular times for you to check on one another?
7. Does it feel like someone took charge and is leading the project? Are you comfortable with this situation?
8. What could you do better as a teammate?
9. What were the biggest challenges you faced in the past two weeks?
10. How do you feel like your performance on the course is going? Do you feel like you have learned valuable knowledge in the past couple weeks?
11. Did you face any problems in the course during this period? If so, is there any situation you believe we need to be aware of?
12. Based on what you have experienced here so far, how do you expect your next couple of weeks here to be? Do you feel excited? Sad? Scared? Please describe.

### Reflection Assignment 1

1. What aspects of C. elegans were surprising to you?
2. What aspects of working in the dry lab were surprising to you?
3. Do you think C. elegans are a practical choice as a model organism for genomics? Explain why or why not?
4. Reflect of a collaborative project were both wet lab and dry lab techniques were utilized. What stood out to you about this collaboration?
5. Following the session, have your perspectives on research changes? If so, in what specific ways?
6. What insights did you gain about Bioinformatics in real research environment?

### Reflection Assignment 2

1. Was the research project an enriching educational experience for you?
2. Did the research project change your perception of the Bioinformatics field? In what way? If not, why?
3. Was the research project adapted to your learning style?
4. One thing you learned that was unexpected.
5. One thing you learned that will probably be very useful in the future.
6. One thing you would have done differently as a teammate.
7. One thing you would have done differently in your research project.

### Peer evaluation

Adapted from: Carver TL, Stickley A: Teamwork in First Year Law Units: Can It Work? Journal of University Teaching & Learning Practice. 2012, 9:1-33.

| **Category** | **4** | **3** | **2** | **1** |
| --- | --- | --- | --- | --- |
| **Responsibility and Engagement** | Performs all assigned duties and does work without being reminded. Attends all meetings on time. | Performs nearly all assigned duties and/or rarely needs reminding. Attends all meetings on time. | Performs few assigned duties and/or often needs reminding. Attends most meetings, sometimes late. | Does not performs assigned duties and relies on others to do the work and/or does not attend meetings. |
| **Quality of Contributions** | Routinely provides useful ideas when participating in the group and in classroom discussion. A definite leader who contributes a lot of effort. Provides work of the highest quality. | Usually provides useful ideas when participating in the group and in classroom discussion. A strong group member who tries hard. Provides high quality work. | Sometimes provides useful ideas when participating in the group and in classroom discussion. A satisfactory group member who does what is required. Provides work that occasionally needs to be checked/redone by other group members to ensure quality. | Rarely provides useful ideas when participating in the group and in classroom discussion. May refuse to participate. Provides work that usually needs to be checked/redone by others to ensure quality. |
| **Time-management** | Routinely uses time well throughout the project to ensure things get done on time. Group does not have to adjust deadlines or work responsibilities because of this person's procrastination. | Usually uses time well throughout the project but may have procrastinated on one thing. Group does not have to adjust deadlines or work responsibilities because of this person's procrastination. | Tends to procrastinate, but always gets things done by the deadlines. Group does not have to adjust deadlines or work responsibilities because of this person's procrastination. | Rarely gets things done by the deadlines AND group has to adjust deadlines or work responsibilities because of this person's inadequate time management. |
| **Cooperation** | Never argue. Provides constructive criticism when appropriate.  Is never publicly dismissive of the project or the work of others. Responds positively to feedback. | Rarely argues unproductively. Sometimes provides constructive criticism when appropriate.  Rarely is publicly dismissive of the project or the work of others. Responds positively to feedback. | Sometimes argues unproductively. Occasionally is publicly dismissive of the project or the work of other members of the group. May not respond positively to feedback. | Usually argues unproductively. Often is publicly dismissive of the project or the work of other members of the group. Does not respond to feedback positively. |
| **Working with Others** | Almost always listens to, shares with, and supports the efforts of others. Tries to keep people working well together. | Usually listens to, shares, with, and supports the efforts of others. Does not cause "waves" in the group. | Often listens to, shares with, and supports the efforts of others, but sometimes is not a good team member. | Rarely listens to, shares with, and supports the efforts of others. Often is not a good team player. |

### An example of PCR gels (MDSC 301 2024)

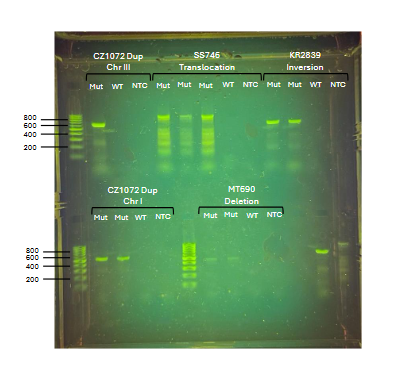

### Post-course survey

1. Which course with a RESEARCH EXPERIENCE did you take this term?

2. I contributed to crafting a question for the RESEARCH EXPERIENCE (e.g. I articulated, created, or identified a research question).

3. The RESEARCH EXPERIENCE contributed to my interest/engagement in the topic.

4. I collected/contributed data in the RESEARCH EXPERIENCE.

5. I analyzed/interpreted data in the RESEARCH EXPERIENCE.

6. I communicated my RESEARCH EXPERIENCE (e.g. I talked about the research experience within my social network to one or more person: such as a classmate, peer, family member, significant other, roommate, mentor, etc...).

7. The RESEARCH EXPERIENCE was directly related to the course topic.

8. The purpose/procedures of the RESEARCH EXPERIENCE were introduced in the syllabus and/or during class activities.

9. The RESEARCH EXPERIENCE was adequately supported by the instruction.

10. The RESEARCH EXPERIENCE was an interesting way to learn about the course material.

11. This RESEARCH EXPERIENCE helped to develop...

· My ability to read and understand scholarly literature primary

· My ability to analyze and synthesize information from academic sources

· My ability to communicate research-based information

· My ability to communicate my ideas

· My ability to work independently

· My ability to collaborate/build relationships/work with others

· My self-confidence

· My resiliency

· My ability to handle challenges

· My ability to navigate ambiguity (ambiguity = unknown factors or outcomes)

· My ability to problem-solve

· My understanding of how academics/scholars/scientists think and work

· My understanding of how practitioners/coaches think and work

· My sense of what careers are available to me

12. My interest in my chosen program of study or degree.

13. My understanding of research: what it is and how it works.

14. I think that participating in a Course-based RESEARCH EXPERIENCE (CURE) is a valuable learning experience.

15. I think that participating in a Course-based RESEARCH EXPERIENCE (CURE) develops students’ academic skills (e.g. one or more of these or related skills: reading and understanding peer reviewed articles, collecting data, critical thinking, problem solving, analysis, collaborating, communicating data and/or findings, working with others, handling problems).

16. I think that participating in a Course-based RESEARCH EXPERIENCE (CURE) helps a student to do well in subsequent classes.

17. I think that participating in a Course-based RESEARCH EXPERIENCE (CURE) contributes to a student’s engagement in the course/program.

18. I think that participating in a Couse-based RESEARCH EXPERIENCE (CURE) clarifies a student’s study/career interests.

19. As a result of this RESEARCH EXPERIENCE, I…

20. This RESEARCH EXPERIENCE helped to develop….

21. Participating in a RESEARCH EXPERIENCE meant….
